## Supplementary figures and images for "Resilience of S309 and AZD7442 monoclonal antibody treatments against infection by SARS-CoV-2 Omicron lineage strains"

### Extended Data Figure 1

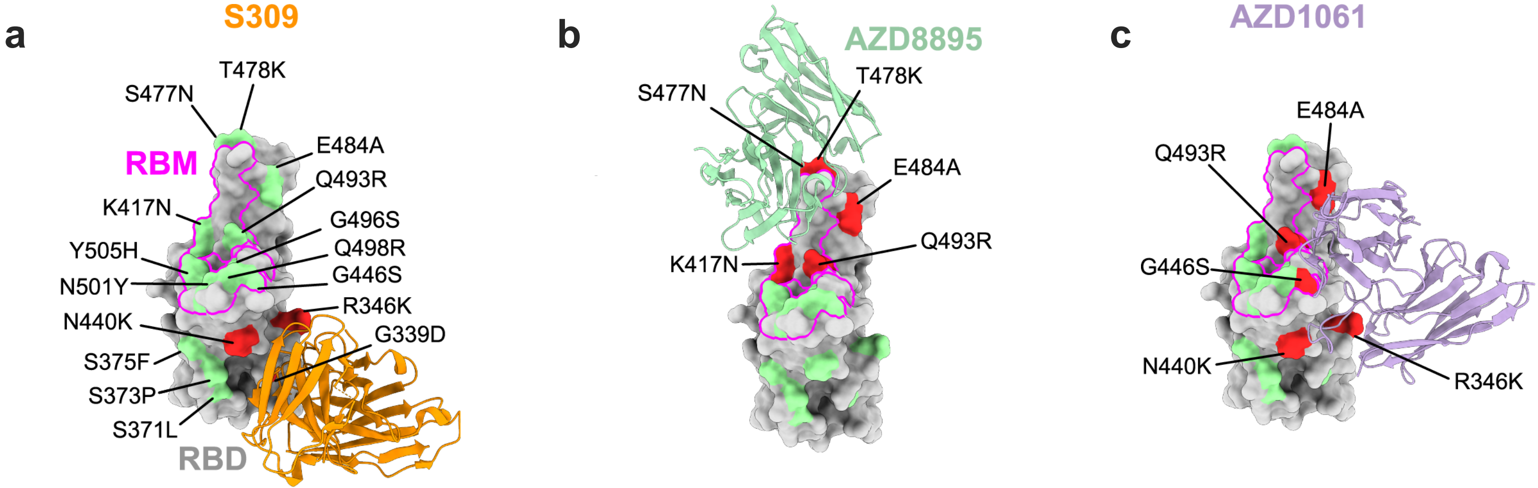


**Extended Data Figure 1**

### Extended Data Figure 2

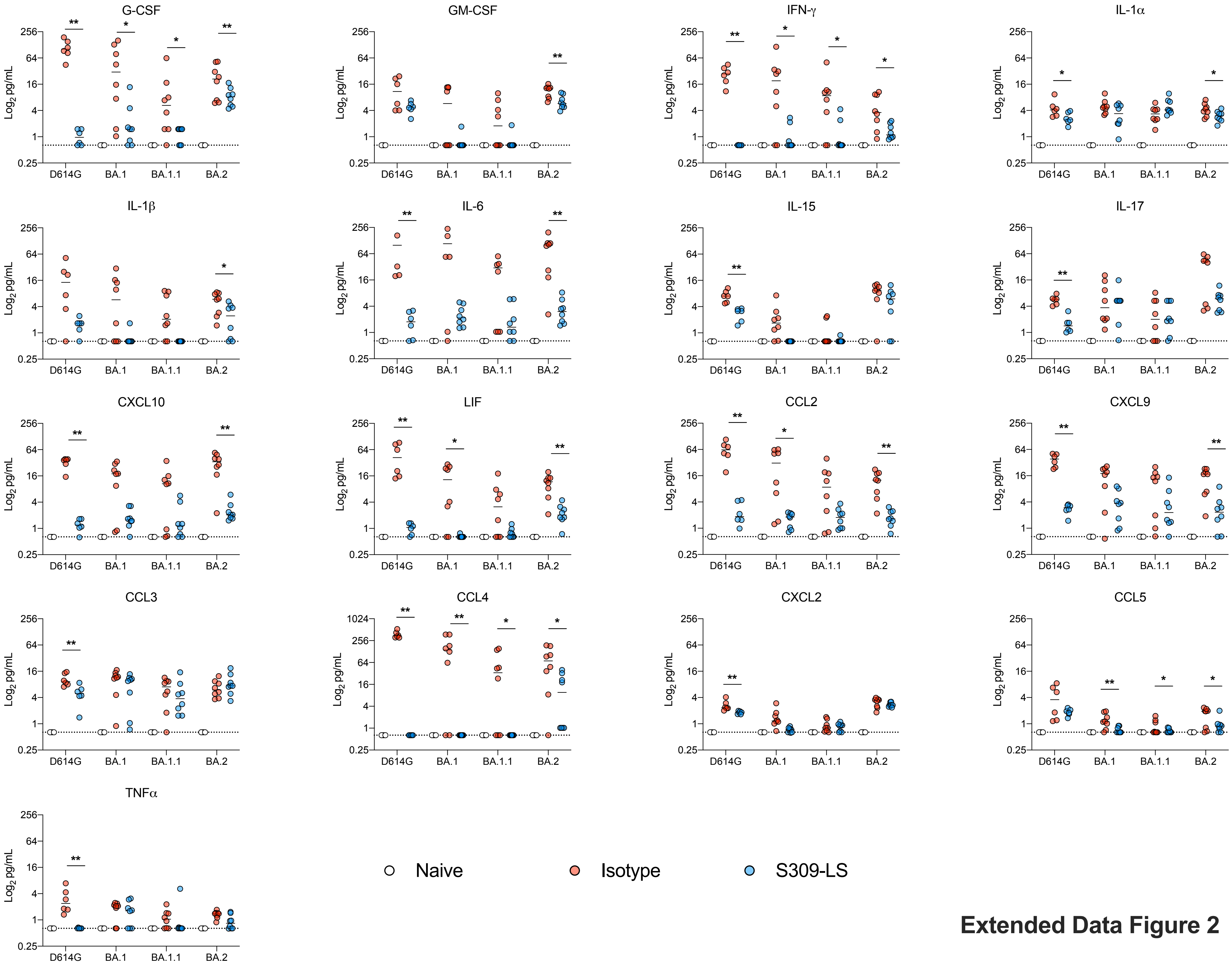


**Extended Data Figure 2**

### Extended Data Figure 3

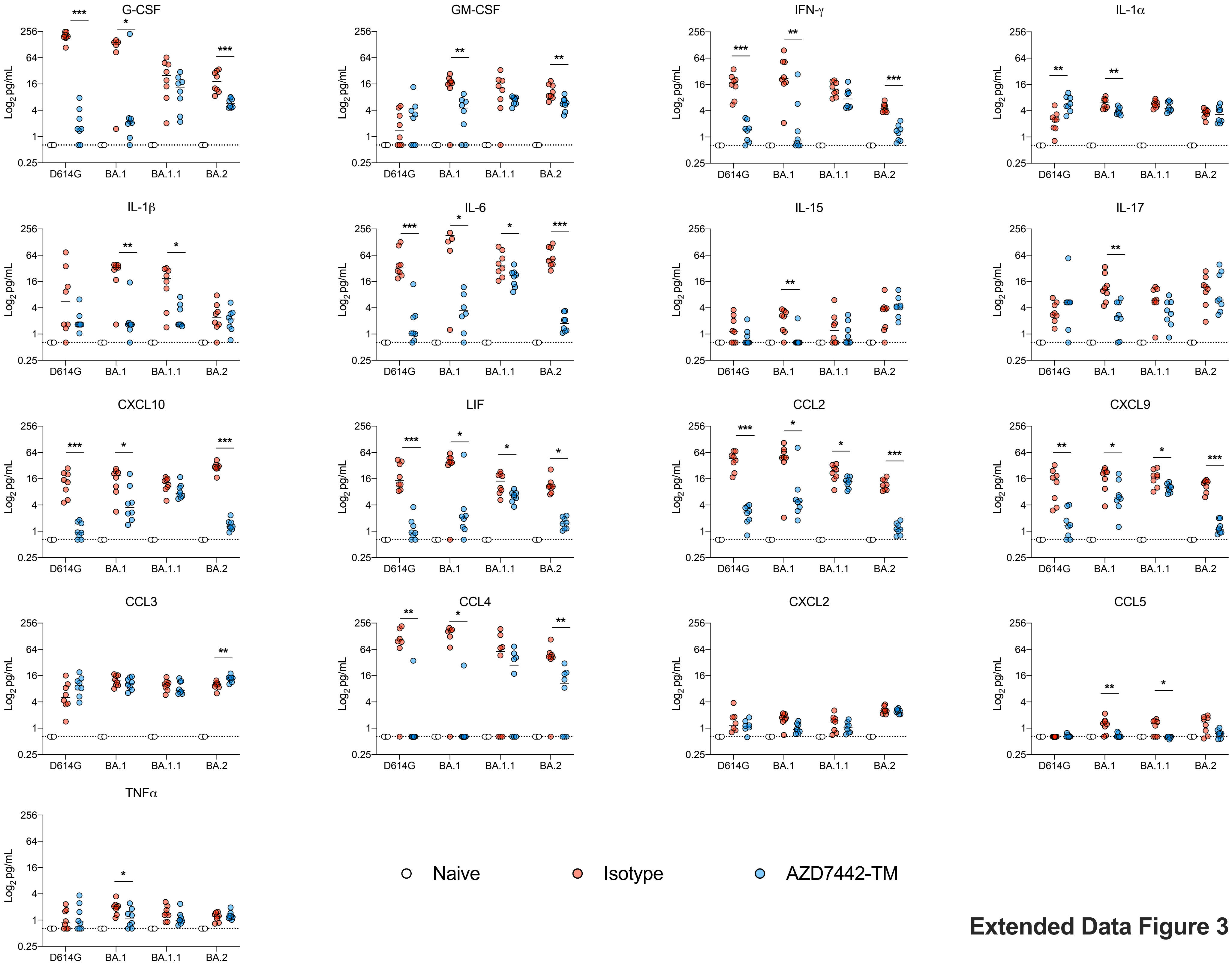


**Extended Data Figure 3**

### Extended Data Figure 4

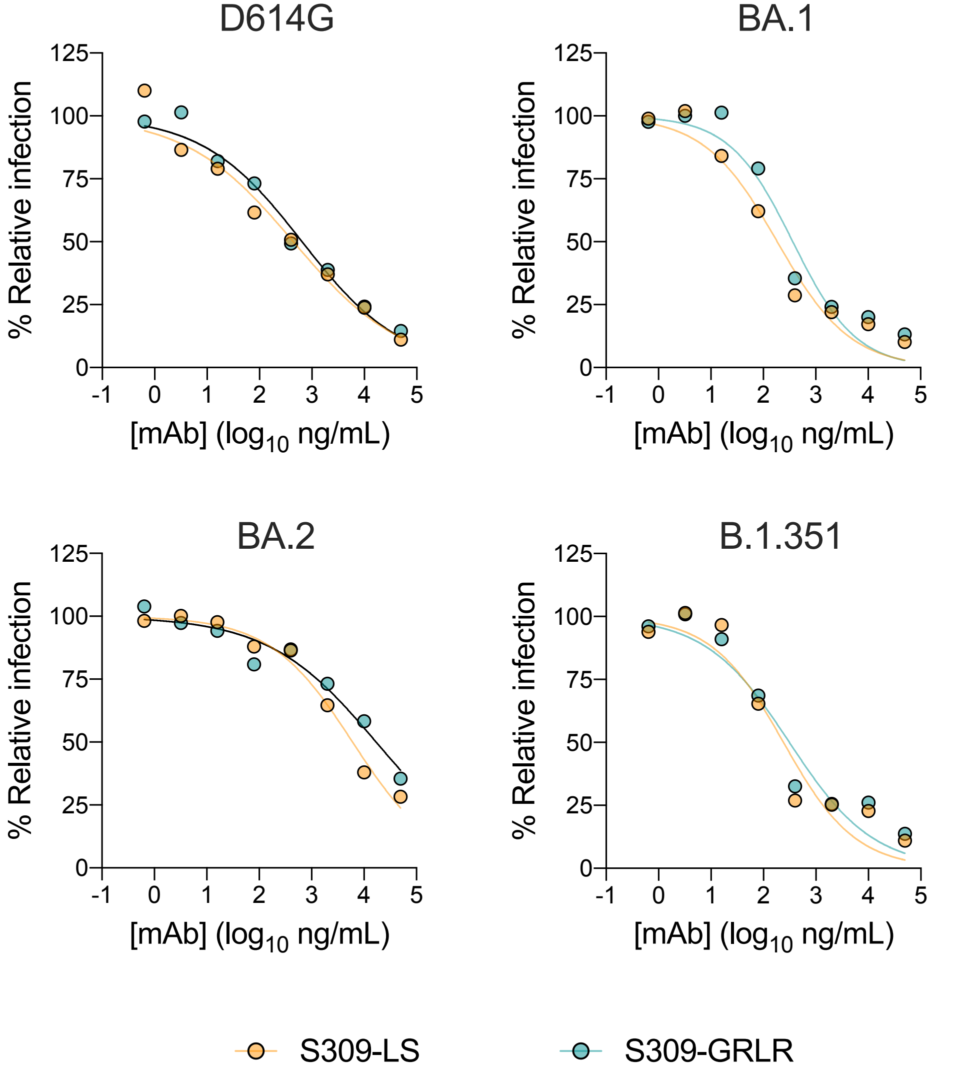


**Extended Data Figure 4**

### Extended Data Figure 5

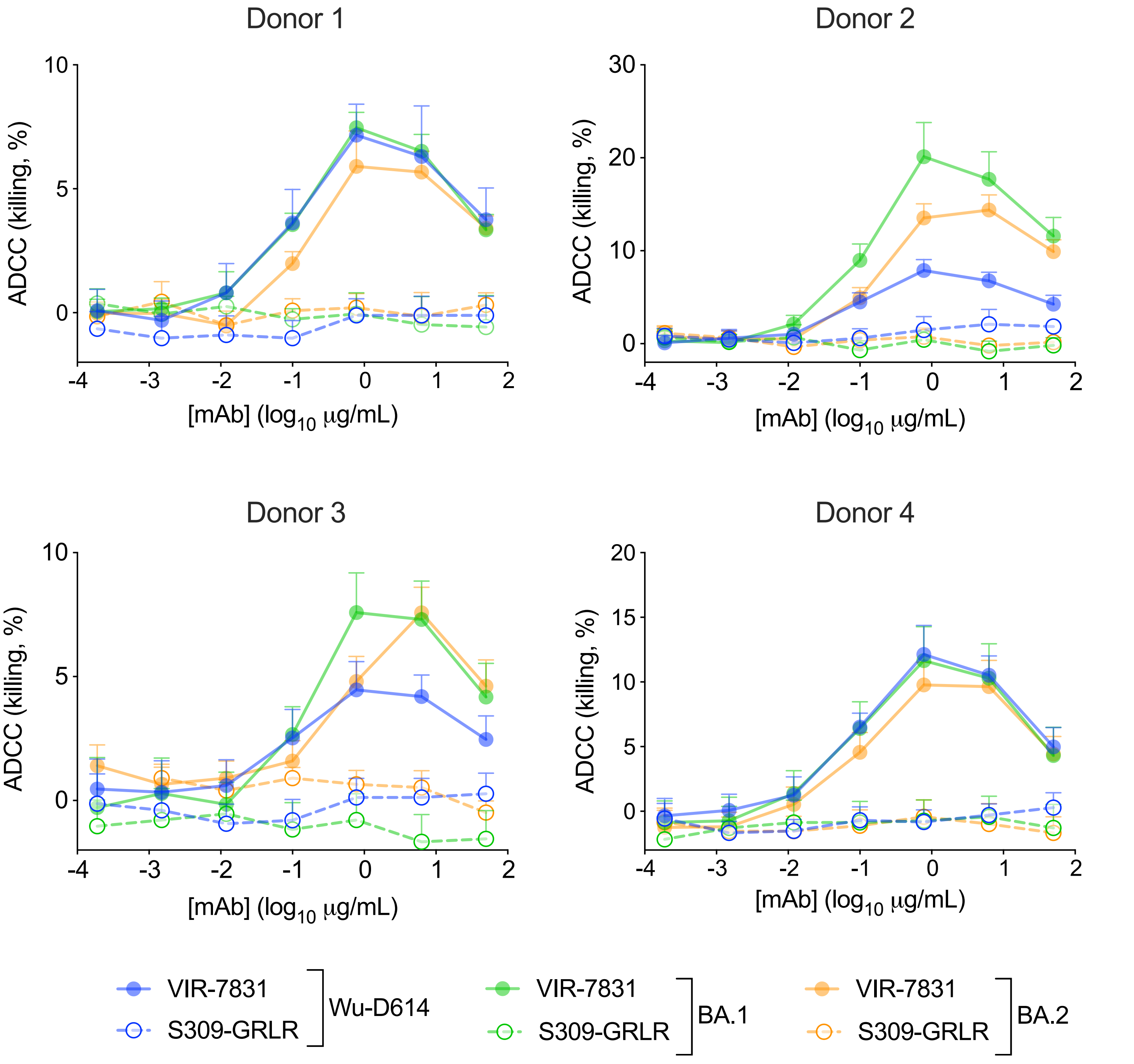


**Extended Data Figure 5**

### Extended Data Figure 6

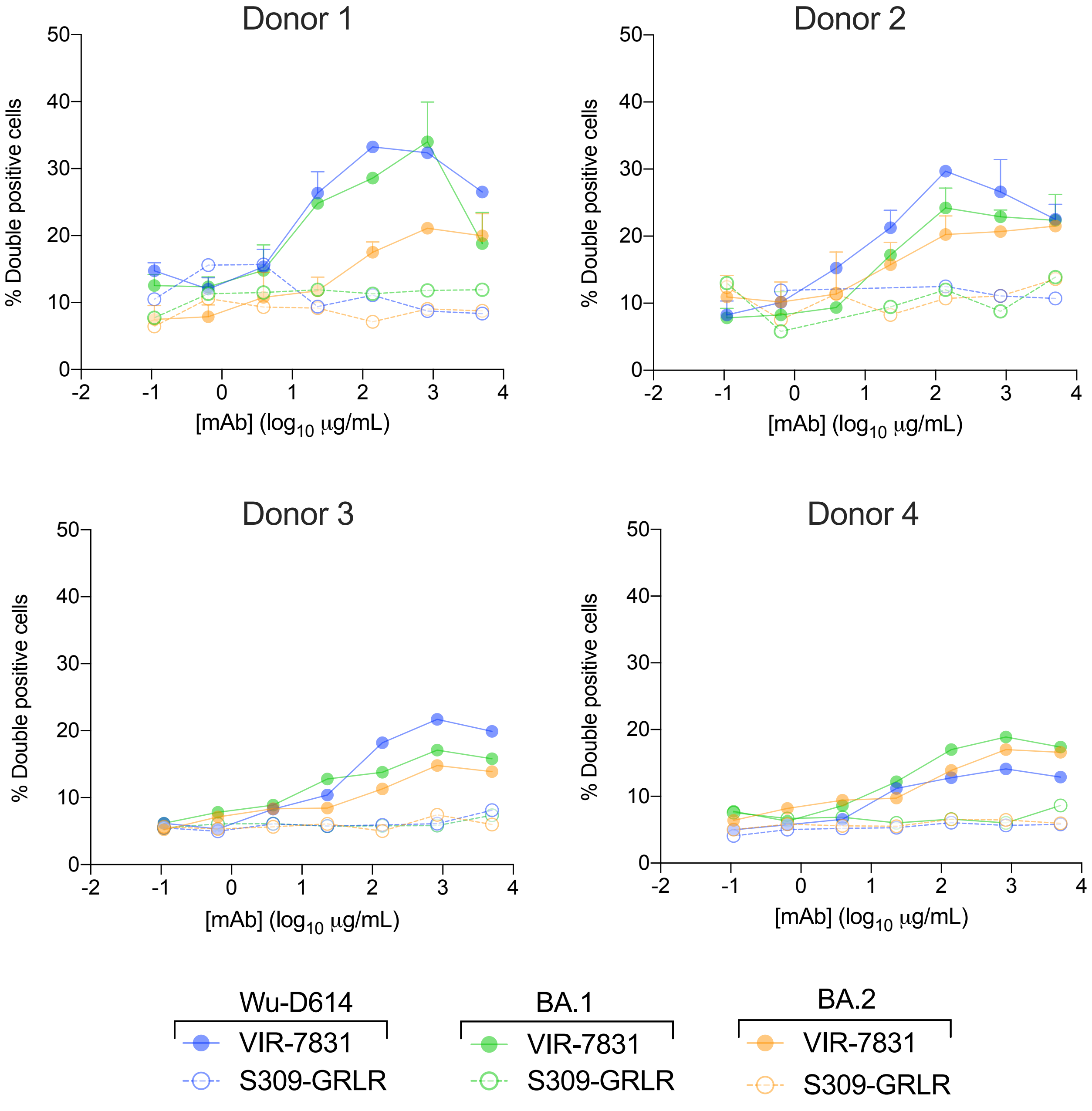


**Extended Data Figure 6**

### Extended Data Figure 7

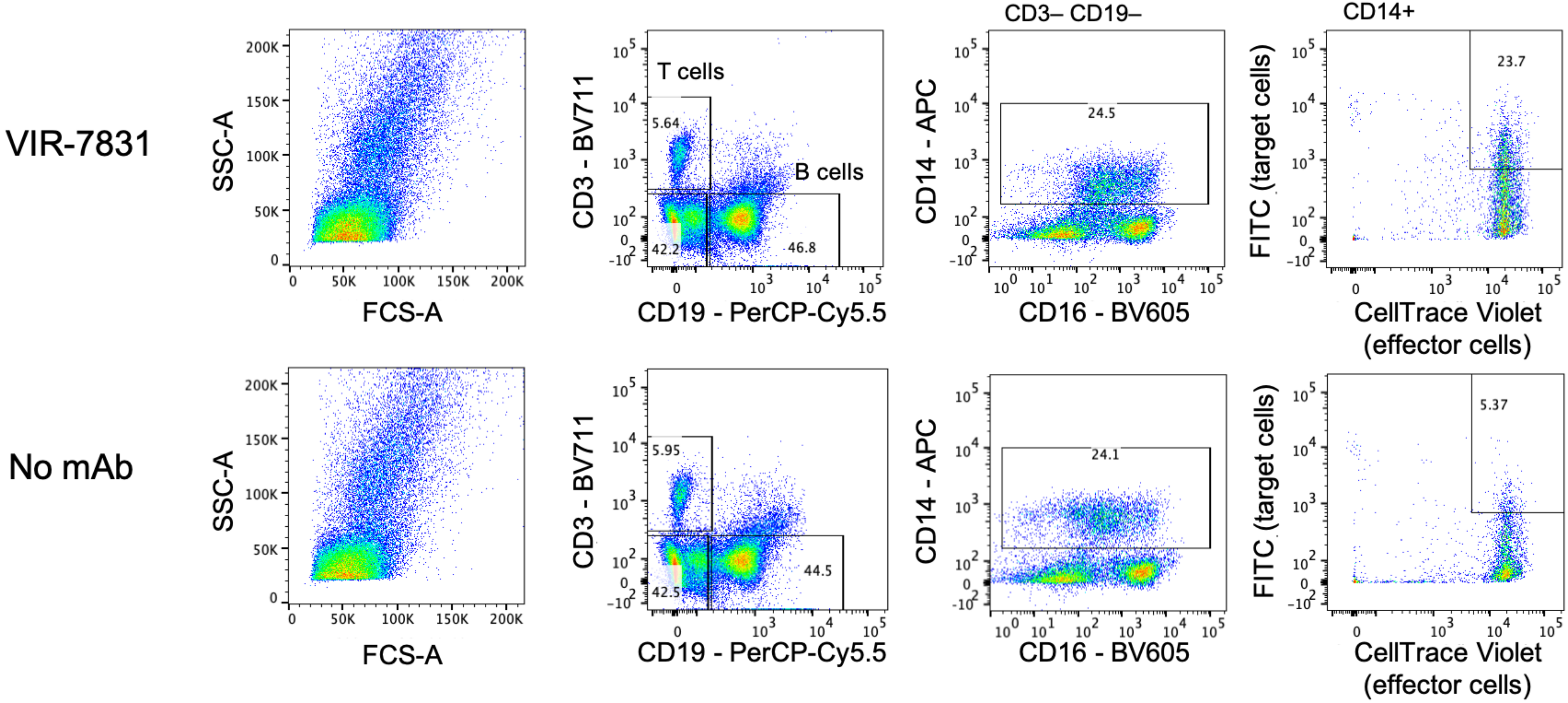


**Extended Data Figure 7**
